# Milk casein prevents inactivation effect of black tea galloylated theaflavins on SARS-CoV-2 *in vitro*

**DOI:** 10.1101/2021.09.27.461930

**Authors:** Eriko Ohgitani, Masaharu Shin-Ya, Masaki Ichitani, Makoto Kobayashi, Takanobu Takihara, Motoki Saito, Hitoshi Kinugasa, Osam Mazda

**Affiliations:** Department of Immunology, Kyoto Prefectural University of Medicine, Kamigyo, Kyoto 602-8566, Japan; Central Research Institute, ITO EN, Ltd., Makinohara, Shizuoka 421-0516, Japan

**Keywords:** novel coronavirus, COVID-19, delta variant, tea, theaflavin, polyphenol, milk, casein

## Abstract

Repeated emergence of highly contagious and potentially immune-evading variant SARS-CoV-2 is posing global health and socioeconomical threats. For suppression of the spread of the virus infection among people, a procedure to inactivate virus in saliva may be useful, because saliva of infected persons is the major origin of droplets and aerosols that mediate viral transmission to nearby persons. We previously reported that SARS-CoV-2 is rapidly and remarkably inactivated by treatment *in vitro* with tea including green tea, roasted green tea, oolong tea and black tea. Tea catechin-derived compounds including theaflavins (TFs) with (a) galloyl moiety(ies) showed this activity. Although black tea is popularly consumed worldwide, a lot of people consume it with sugar, milk, lemon juice, and so on. But it has not been determined whether these ingredients may influence the inactivation effect of black tea against SARS-CoV-2. Moreover, it has not been revealed whether black tea is capable of inactivating variant viruses such as delta variant. Here we examined the effect of black tea on some variants in the presence or absence of sugar, milk, and lemon juice *in vitro*. Black tea and galloylated TFs remarkably inactivated alpha, gamma, delta and kappa variants. Intriguingly, an addition of milk but not sugar and lemon juice totally prevented black tea from inactivating alpha and delta variant viruses. The suppressive effect was also exerted by milk casein. These results suggest the possibility that intake of black tea without milk by infected persons may result in inactivation of the virus in saliva and attenuation of spread of SARS-CoV-2 to nearby persons through droplets. Clinical studies are required to investigate this possibility.

## 1. Introduction

SARS-CoV-2 is mainly transmitted by droplets and aerosols that are derived from saliva of COVID-19 patients and asymptomatic infected persons, although bronchial and laryngeal aerosols may also be involved in the viral transmission [1-5]. Saliva containing the virus is scattered by speaking, coughing and sneezing from the oral cavity to environment and form the droplets and aerosols that could reach nasal and oral mucosa of nearby persons to cause infection. We considered that inactivation of virus in saliva may result in attenuation of viral spread among people, and explored various food ingredients that inactivate the SARS-CoV-2. We reported that tea including green tea, roasted green tea, oolong tea and black tea significantly reduced infectivity of conventional SARS-CoV-2 *in vitro* [6,7]. We also found that (-) epigallocatechin-gallate (EGCG), a tea catechin contained in tea leaves and green tea at high concentrations, rapidly and powerfully inactivated the virus, while stronger effects were exerted by black tea ingredients, theaflavins (TFs) with (a) 3 and/or 3’ galloyl moiety(ies) (theaflavin-3-gallate (TF3G), theaflavin-3’-gallate (TF3’G), and theaflavin-3,3’-O-digallate (TFDG)) and theasinensin A (TSA) that are produced from catechins by enzymatic oxidization during manufacturing of black tea [7].

More recently, variant SARS-CoV-2 viruses that are highly contagious and may evade immunity have emerged and spread worldwide [8-10]. But it has not been clarified whether tea ingredients also inactivate these variants.

Meanwhile, more than 70% of tea is manufactured into black tea that is consumed all over the world. People in some countries/areas consume larger amounts of black tea *per capita* than those in other countries/areas, but any epidemiological analysis have not demonstrated a significant negative correlation between black tea consumption and infected population. Black tea is often consumed with sugar, milk, lemon juice, honey, jam, cinnamon, etc. These ingredients could potentially influence the activity of black tea to block the virus infection.

In this study we examined whether black tea inactivated the variants of the SARS-CoV-2 *in vitro*. We also assessed whether an addition of sugar, milk, or lemon juice influenced the anti-virus effect of black tea.

## 2. Results

### 2.1. Black Tea Inactivated Alpha, Gamma, Delta and Kappa Variants of SARS-CoV-2

Conventional and variants of SARS-CoV-2 (Supplemental Table 1) were treated with black tea, and the titers of the viruses were determined (Fig. 1a). The alpha and gamma variants were as remarkably inactivated by black tea as conventional SARS-CoV-2 (Fig. 1b). Similar results were also obtained with delta and kappa variants (data not shown).

**Fig. 1.**
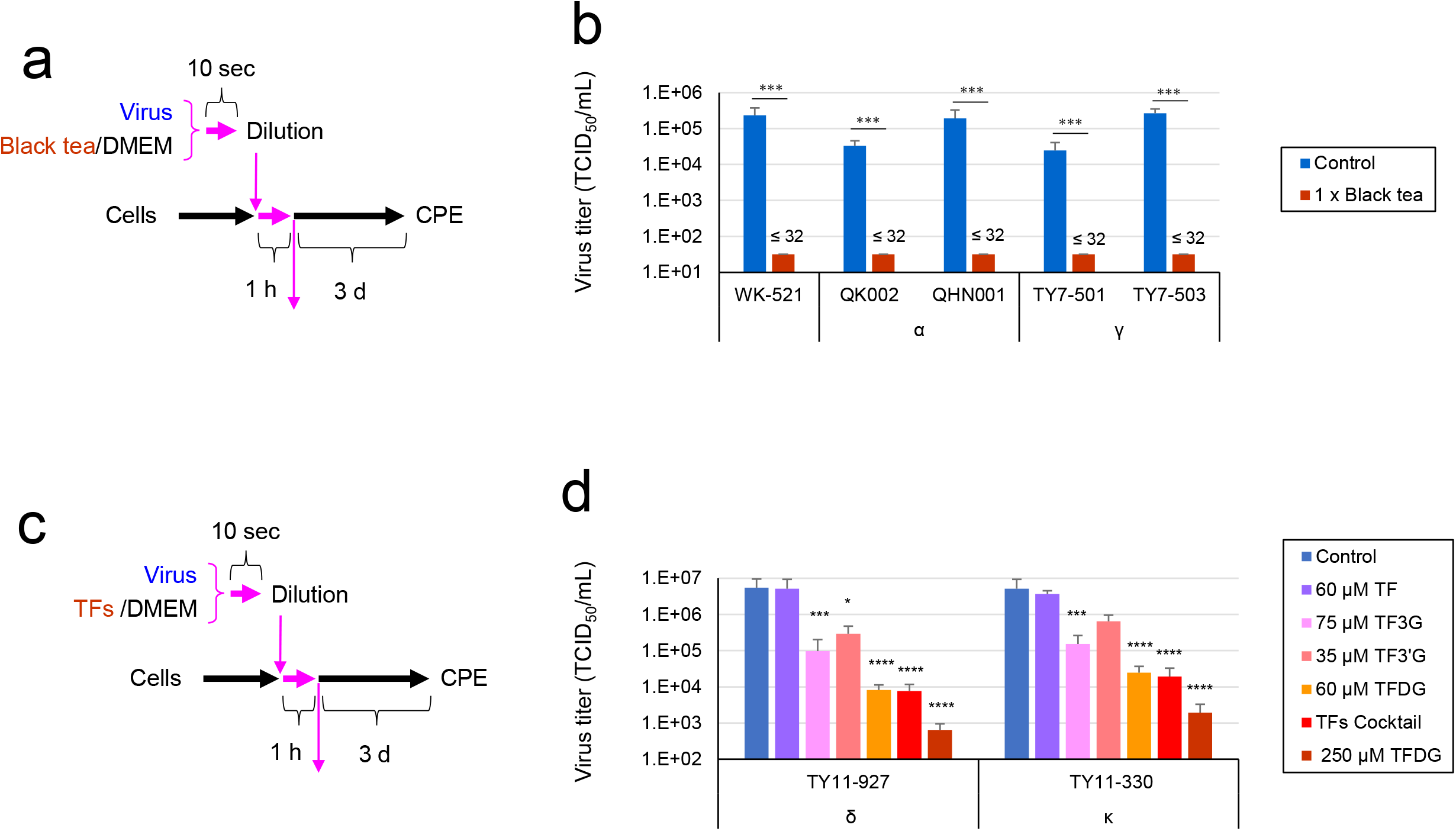
Titration of SARS-CoV-2 variants treated with black tea or TFs. (a and b) Indicated viruses were treated with x1 black tea/DMEM or DMEM for 10 sec, immediately followed by a 10-fold serial dilution and infection to VeroE6/TMPRSS2 cells to measure TCID_50_ values as described in the Materials and Methods. Scheme of experiments (a), and virus titer of each sample (means ± S.D., n = 3) (b) are shown. (c and d) Viruses were treated with the indicated concentrations of TFs/DMEM solution for 10 sec, immediately followed by a 10-fold serial dilution and infection to VeroE6/TMPRSS2 cells as above. Scheme of experiments (c), and virus titer of each sample (means ± S.D., n = 3) (d) are shown. * p < 0.05, ** p < 0.01 and *** p < 0.001 vs. Control (DW) by Tukey’s multiple comparison test.

Effects of TF and its gallate esters on SARS-CoV-2 were examined. The black tea that we used in the experiments contains approximately 60 µM TF, 75 µM TF3G, 35 µM TF3’G, and 60 µM TFDG [7]. These concentrations of TFs were tested for the inhibitory activity against delta and kappa variants (Fig. 1c). Another aliquot of virus was treated with a cocktail of these TFs. As shown Fig. 1 d, non-galloylated TF at 60 µM failed to reduce the virus titers, whereas both viruses were remarkably inactivated by 75 µM TF3G. The virus titers more remarkably fell after treatment with 60 µM TFDG, and to a similar extent, with the cocktail of TFs. We previously compared effects of the same concentration of these four TFs, and found that TFDG most strongly inhibited conventional SARS-CoV-2, while TF3G and TF3’G showed lower activities, and any significant inhibitory effect was not exhibited by non-galloylated TF [7]. TFDG also inactivated the alpha and gamma variants (data not shown). The present data on the delta and kappa variants are consistent with the previous findings on conventional virus, and TFDG most principally contributed to the anti-virus effect of black tea among the TFs tested.

### 2.2. Black Tea Inactivated the Delta Variant that Were Diluted in Human Saliva

To estimate the possibility that tea consumption by infected persons results in inactivation of the virus in saliva, the delta variant virus was diluted in saliva from four healthy donors (Supplemental Table 2), and the mixture was treated with black tea at 1:1 (vol:vol) (Fig. 2a). After infection, viral RNA replication in the cells was evaluated by real time RT-PCR analysis. As shown in Fig. 2b, viral RNA was significantly fewer in the cells infected with black tea-treated virus than in the cells infected with non-treated virus. We also quantified viral RNA in the culture supernatant to evaluate secondary virus released from the infected cells. It was revealed that the treatment of the virus with black tea resulted in a significantly lower degree of secondary virus production from the infected cells.

**Fig. 2.**
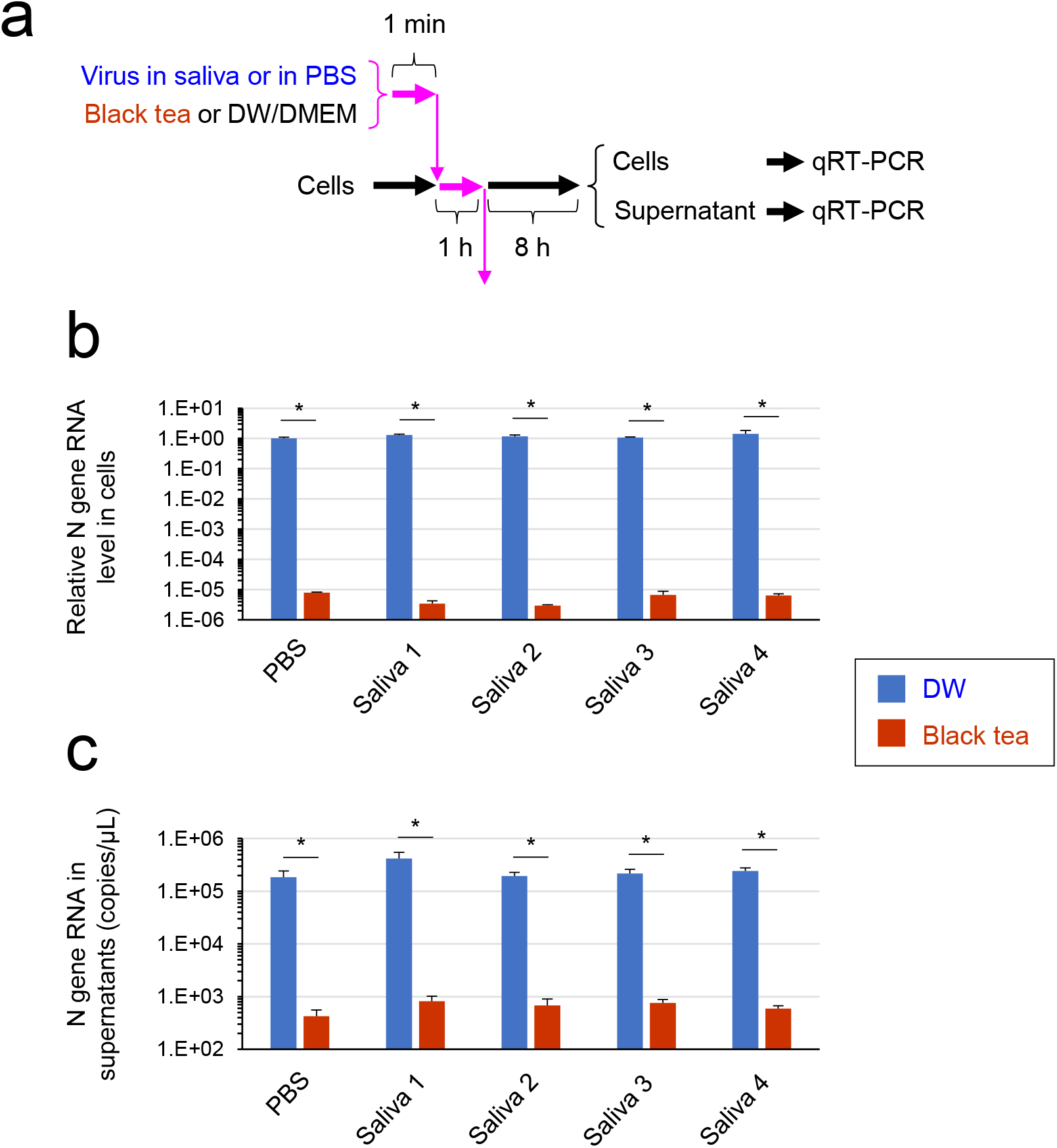
Black tea inactivated delta variant of SARS-CoV-2 diluted in human saliva. SARS-CoV-2 was diluted in saliva from four healthy donors (Numbers 1-4 in Supplementary Table S2) or in distilled water (DW) as a control, and treated with x1 black tea or DW as described in the Materials and Methods. One min later the mixture was added to the cells for 1 h, followed by replacement of the supernatant by fresh culture medium and subsequent culture for 8 h. RNA was extracted from the cells and the culture supernatants, and real time-RT-PCR was performed to evaluate viral N gene RNA levels. Scheme of experiments (a) and means ± S.D. of RNA levels (N = 3) (b and c) are shown. * p < 0.05, between groups.

### 2.3. Milk Casein Prevented Inactivation of SARS-CoV-2 by Black Tea

We supplemented black tea with sugar, milk and/or lemon juice at ratios comparable to ordinary beverages (6 g sugar, 40 mL milk, and 2 mL lemon juice per 160 mL of tea), and examined whether the supplementation influenced the ability of black tea to inactivate the alpha and delta variants diluted in human saliva (Fig. 3a). Although black tea significantly inactivated the SARS-CoV-2 variants, the anti-virus activity of the black tea was totally hampered by an addition of milk (Fig. 3b-c). Neither sugar nor lemon juice significantly affected the titers of the virus.

**Fig. 3.**
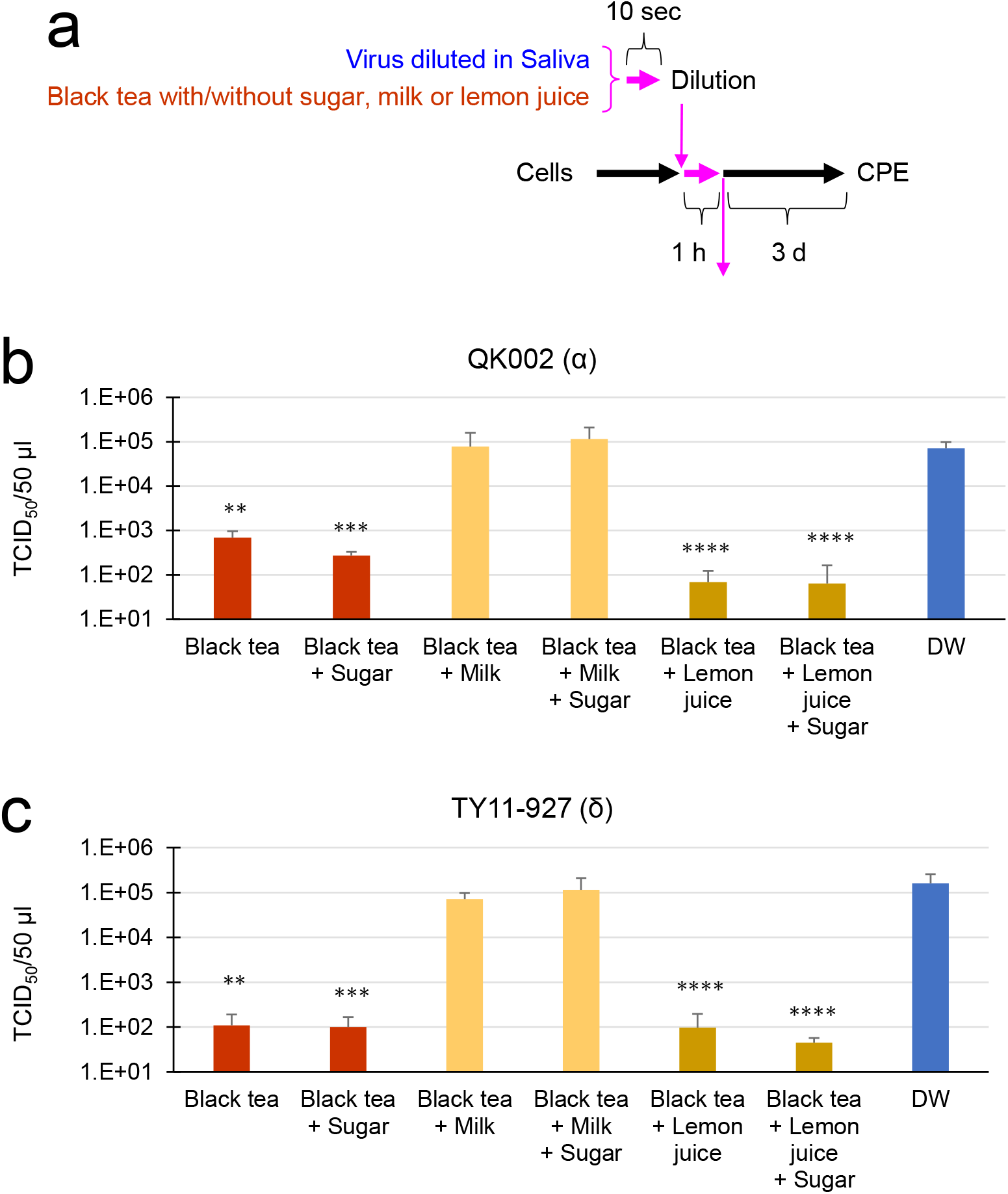
An addition of milk cancelled anti-SARS-CoV-2 effect of black tea. a) Scheme of experiments are shown. b and c) QK002 (alpha variant)(b) and TY11-927 (delta variant)(c) was diluted in saliva from healthy human donors (Numbers 5 and 6 in Supplementary Table S2 were used in (b) and (c), respectively). Black tea supplemented with milk, sugar, and/or lemon juice was added to the virus/saliva for 10 sec as described in the Materials and Methods, immediately followed by a serial dilution with MS. TCID_50_ assay was performed as described in the Materials and Methods. Virus titer of each sample (means ± S.D.) is shown (n = 3). * p < 0.05, ** p < 0.01 and *** p < 0.001 vs. Control (DW) by Tukey’s multiple comparison test.

What is the active component in milk that inhibits the anti-SARS-CoV-2 activity of black tea? Some previous literatures reported that caseins form a complex with tea catechins [11-14]. Because theaflavins with (a) gallate ester(s) are derived from, and share common structures with, catechins, we examined possible involvement of casein in the inhibitory activity of milk. We prepared a casein solution at a concentration of 2.7% (w/v) that is comparable to a standard concentration of casein in ordinary cow’s milk (Fig. 4a). For an example, an addition of 25% (vol) of this casein solution to black tea (thus, 250 µL casein solution was added into 1 mL of tea) simulates an addition of 25 mL of milk to 100 mL of black tea, resulting in an approximate casein concentration of 0.54% (w/v). As results, casein dose-dependently prevented black tea from reducing the virus titer (Fig. 4b-c).

**Fig. 4.**
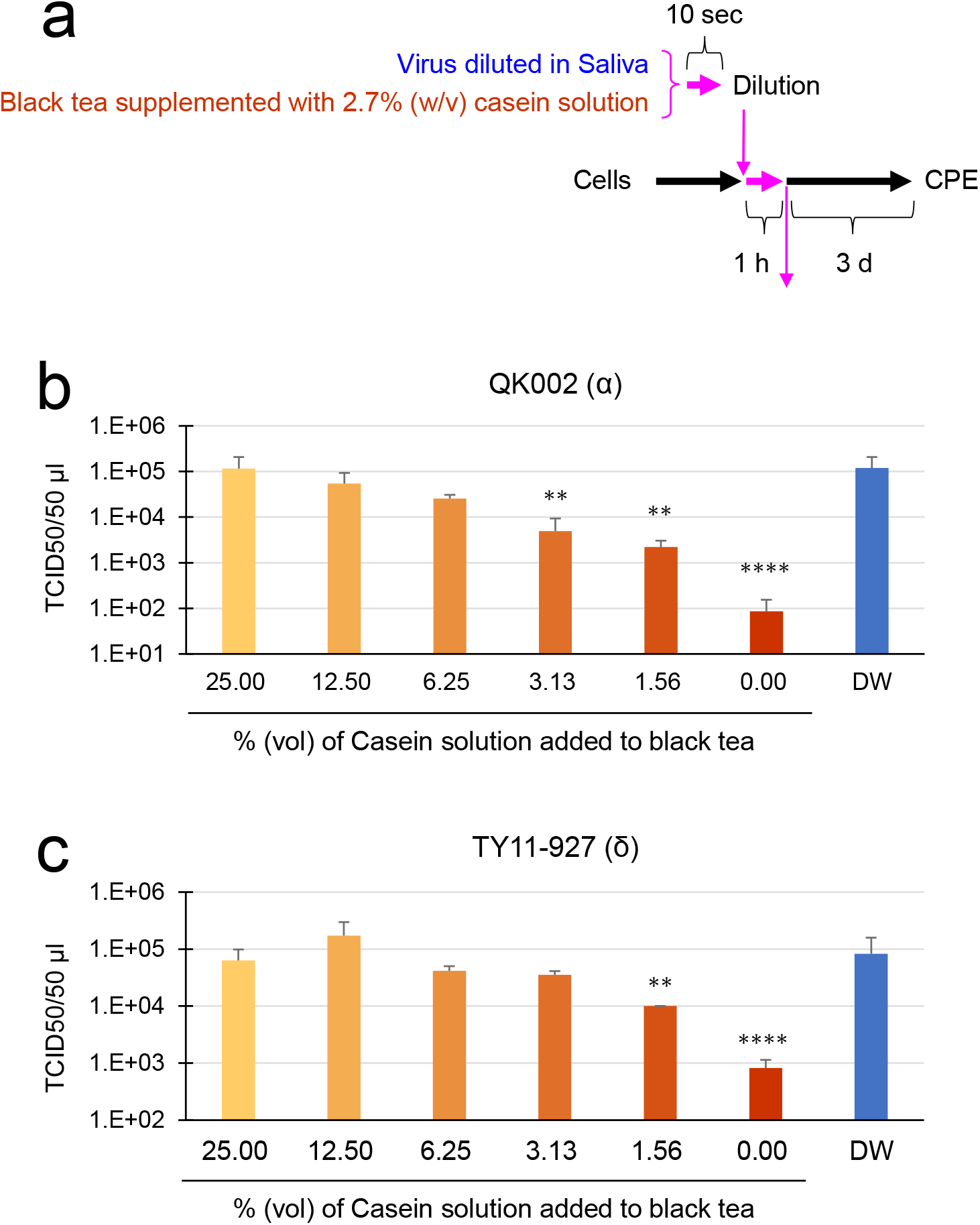
An addition of milk casein solution cancelled anti-SARS-CoV-2 effect of black tea. a) Scheme of experiments are shown. b and c) 2.7% casein solution was added to black tea at the indicated % (vol)(vol of black tea is regarded as 100%). QK002 (b) and TY11-927 (c) was diluted in saliva from healthy human donors (Numbers 7 and 8 in Supplementary Table S2 were used in (b) and (c), respectively), and mixed with the back tea with the casein solution (vol:vol=1:1) for 10 sec. Immediately the virus/saliva/tea mixture was serially diluted with MS. TCID_50_ assay was performed as described in the Materials and Methods. Virus titer of each sample (means ± S.D.) is shown (n = 3). * p < 0.05, ** p < 0.01 and *** p < 0.001 vs. Control (DW) by Tukey’s multiple comparison test.

## 3. Discussion

In this study we demonstrated that black tea and its constituent galloylated TFs inactivate variant SARS-CoV-2 viruses including the delta variant *in vitro*. The inactivation effect was counteracted by an addition of milk casein into the black tea.

Bovine milk contains 32-37 g/L of proteins, and caseins account for about 80% of the whole milk proteins [15,16]. Caseins consist of four polypeptides, i.e., alpha_S1_-, alpha_S2_-, beta- and kappa-caseins, and form large colloidal particles, the casein micelles, that are dispersed in milk [15-17]. Caseins are involved in transport of calcium and phosphorus. Previous reports demonstrated that caseins bound to catechins including EGCG [13,14,18,19], leading to a significant decrease in antioxidant activity of catechins and non-galloylated TF [13,19]. Addition of milk also blocked the vascular protection effect of black tea [18]. Our results suggest that galloylated TFs may also bind to caseins resulting in a loss of anti-SARS-CoV-2 effect of black tea (Fig. 4).

Polyphenols including TFs are involved in flavor, bitter taste and astringency of tea [11,12]. Addition of milk reduces the bitterness and astringency of tea, probably due to the binding of caseins to catechins and TFs.

Tea drinkers add various amount of milk to their black tea infusions. In our study, additions of 6.25% and 3.31% (vol) of 2.7% casein solution into black tea totally blocked the inactivation effect against alpha and delta variants, respectively (Fig. 4b and c). This means that approximately 4 and 8 mL of milk completely block the anti-virus effect of a cup of tea (assuming its volume to be 125 mL) against delta and alpha variants, respectively. A smaller volume of milk partially reduced the anti-virus activities. But the relationship between the ratios of milk vs. tea infusions and degrees of counteraction of anti-SARS-CoV-2 effect may differ depending on the types of tea and milk.

The alpha, gamma, delta and kappa variants of SARS-CoV-2 have N501Y, N501Y plus E484K, L452R, L452R plus E484Q mutations, respectively, in the viral Spike proteins [8-10,20,21]. These amino acid substitutions may be associated with the high contagiousness and/or immune escape of the variants [20,21]. We previously showed that TFDG may bind to the receptor binding domain (RBD) of the Spike proteins [7]. The present data suggest that the amino acid substitutions in these variant viruses may not crucially affect the interaction between galloylated TFs to RBD.

The present findings that black tea inactivates SARS-CoV-2 only in the absence of milk may partly explain the reason why the infected population in a country/area is not negatively correlated with black tea consumption in the country/area. Various other factors including geographical, genetical, cultural, and social ones also influence the degree of virus spread among particular population.

Ingestion of tea at appropriate occasions may potentially attenuate viral spread to nearby persons, as we previously discussed [6,7], and black tea without milk may be applicable to this purpose. Besides, repetitive ingestion of milk-free black tea shortly after SARS-CoV-2 infection could potentially bring a beneficial effect on disease progression in infected persons, because viral amplification in the oral cavity at the earlies stages of infection may play important roles in subsequent viral infection in the lung [22]. Clinical studies are required to investigate these possibilities.

## Acknowledgments

We thank Japan National Institute of Infectious Diseases (Tokyo, Japan) for kindly providing us with the viruses.

## 4. Materials and Methods

### 4.1. Virus, Cells, and Culture Medium

The SARS-CoV-2 shown in Supplementary Table S1 were kindly provided from Japan National Institute of Infectious Diseases (Tokyo, Japan) and propagated using VeroE6/TMPRSS2 cells [23] that were obtained from Japanese Collection of Research Biosources Cell Bank, National Institute of Biomedical Innovation (Osaka, Japan). Cells were cultured in DMEM supplemented with G418 disulfate (1 mg/mL), penicillin (100 units/mL), streptomycin (100 μg/mL), 5% fetal bovine serum at 37 °C in a 5% CO_2_/95% humidified atmosphere (standard conditions).

### 4.2. Reagents

Black tea infusions were prepared by soaking 40 g of ground up and homogenized tea leaves in 2,000 mL water at 80 °C for 30 min. After centrifugation at 4,000 rpm for 15 min, supernatants were collected and filtrated through Toyo No. 2 filter papers, followed by evaporation and freeze-drying. TF, TF3G, TF3’G, and TFDG were purchased from FUJIFILM Wako Pure Chemical Corporation (Osaka, Japan). Milk, sugar and lemon were purchased at a grocery store in Kyoto. Casein from milk was purchased from FUJIFILM Wako Pure Chemical Corporation (Osaka, Japan). Hammarsten bovine casein was purchased from Sigma-Aldrich. Human saliva was purchased from Lee Biosolutions (Maryland Heights, MO, USA) (Supplementary Table S2) [6].

### 4.3. TCID_50_ Assay for Virus Pretreated with Tea

Freeze-dried powders of black tea were dissolved in sterilized distilled water at 78 °C to prepare x 2 concentration of original tea. After chilling at room temperature, each solution was passed through a 0.45 μm filter, and placed into 96-well-plates at 50 μL/well (N=3). SARS-CoV-2 suspension was added to the plate at 5 × 10^5^ TCID_50_/50 μL/well, and 10 sec later the virus/tea mixture was serially diluted 10-fold with MS. Chilled on ice, 50 μL of each sample was added to the VeroE6/TMPRSS2 cells that had been seeded into 96-well-plates at 5 × 10^4^/100 μL/well a day before (N=4). Cells were incubated for 1 h, followed by replacement of the supernatant by fresh 100 μL MS. After culture for 3 days, cells were washed, fixed, and stained with crystal violet solution to estimate CPE as described [24].

### 4.4. Calculation of TCID_50_ Values

TCID_50_ values were calculated by Reed–Muench method as described elsewhere. If one or more triplicate wells of the lowest dilution of a sample did not show CPE, the highest possible average of TCID_50_ value was calculated for the sample.

### 4.5. TCID_50_ Assay for Virus Pretreated with TFs

Solutions of TF, TF3G, TF3’G, and TFDG were diluted in DMEM, and SARS-CoV-2 suspension (5.0 × 10^5^ TCID_50_/50 μL) was added followed by incubation at room temperature for 10 sec. Immediately, the mixture was serially diluted 10-fold with MS, and TCID_50_ assay and calculation of TCID_50_ values were performed as above.

### 4.6. TCID_50_ Assay for Virus that Was Diluted in Saliva and Treated with Black Tea with Milk, Sugar, Lemon Juice and/or Casein

Freeze-dried powders of black tea extract were dissolved in sterilized distilled water at 78 °C to prepare x 1 concentration of original tea. After chilling at room temperature, each solution was supplemented with sugar (6 g per 160 mL of tea), milk (40 mL per 160 mL of tea), and/or lemon juice (2 mL per 160 mL of tea). In some experiments, each tea solution was supplemented with various doses of 2.7% (w/v) casein solution. Each sample was passed through a 0.45 μm filter. Human saliva was sterilized by UV irradiation for 30 min. Virus suspension (3.0 × 10^5^ TCID_50_/5 μL) was mixed with 45 μL of saliva or water, followed by an addition of tea at 1:1 (vol:vol) for 10 sec. Immediately, the virus/saliva/tea mixture was serially diluted at 10-fold with MS in 96-well-plates. Chilled on ice, 100 μL of each sample was added to the VeroE6/TMPRSS2 cells and TCID_50_ assay was performed as above.

### 4.7. Real Time-RT-PCR for Virus that Was Diluted in Saliva and Treated with Black Tea

Freeze-dried powders of black tea were dissolved in distilled water at 78 °C to prepare x4 concentration of original tea. After chilling at room temperature, each solution was passed through a 0.45 μm filter, and mixed with the same volume of x2 serum-free DMEM. Each sample was added to the same volume of SARS-CoV-2 diluted in UV-irradiated human saliva (6.3 × 10^6^ TCID_50_/mL) as above. After incubated at room temperature for 1 min, the virus/tea/saliva mixture was added to the VeroE6/TMPRSS2 cells that had been seeded into 24-well-plates at 2.5 × 10^5^/well a day before. After culture for 8 h under the standard conditions, cells and culture supernatant were harvested, and RNA was extracted using TRI Reagent^(C)^ LS (Molecular Research Center, Inc., Montgomery Road, Cincinnati, OH, USA). After reverse-transcription using ReverTra Ace^(C)^ qPCR RT Master Mix (Toyobo, Shiga, Japan), quantitative real-time PCR was performed using a Step-One Plus Real-Time PCR system (Applied Biosystems, Foster City, CA, USA) and the following set of primers/probes specific for viral N gene: Forward primer, 5’-AAATTTTGGGGACCAGGAAC-3’; reverse primer, 5’-TGG-CAGCTGTGTAGGTCAAC-3’; and probe, 5’-(FAM) ATGTCGCGCATTGGCATGGA (BHQ)-3’. Ct value for each sample was calculated by StepOne Software (ABI, Warrington, UK). N gene RNA levels in cells were normalized with respect to the 18S rRNA level in each sample, and expressed relative to the value for the control that was infected with untreated-virus (set to 1.0). N gene RNA levels in supernatants were expressed as copy numbers per µL.

### 4.8. Statistical Analysis

Statistical significance was analyzed by Tukey’s multiple comparison test (Figures 1d, 3b, 3c, 4b, and 4c) and Student’s *t* test (Figures 1b, 2b and 2c). *p* < 0.05 was considered significant.

---

**Supplementary Table 1.**
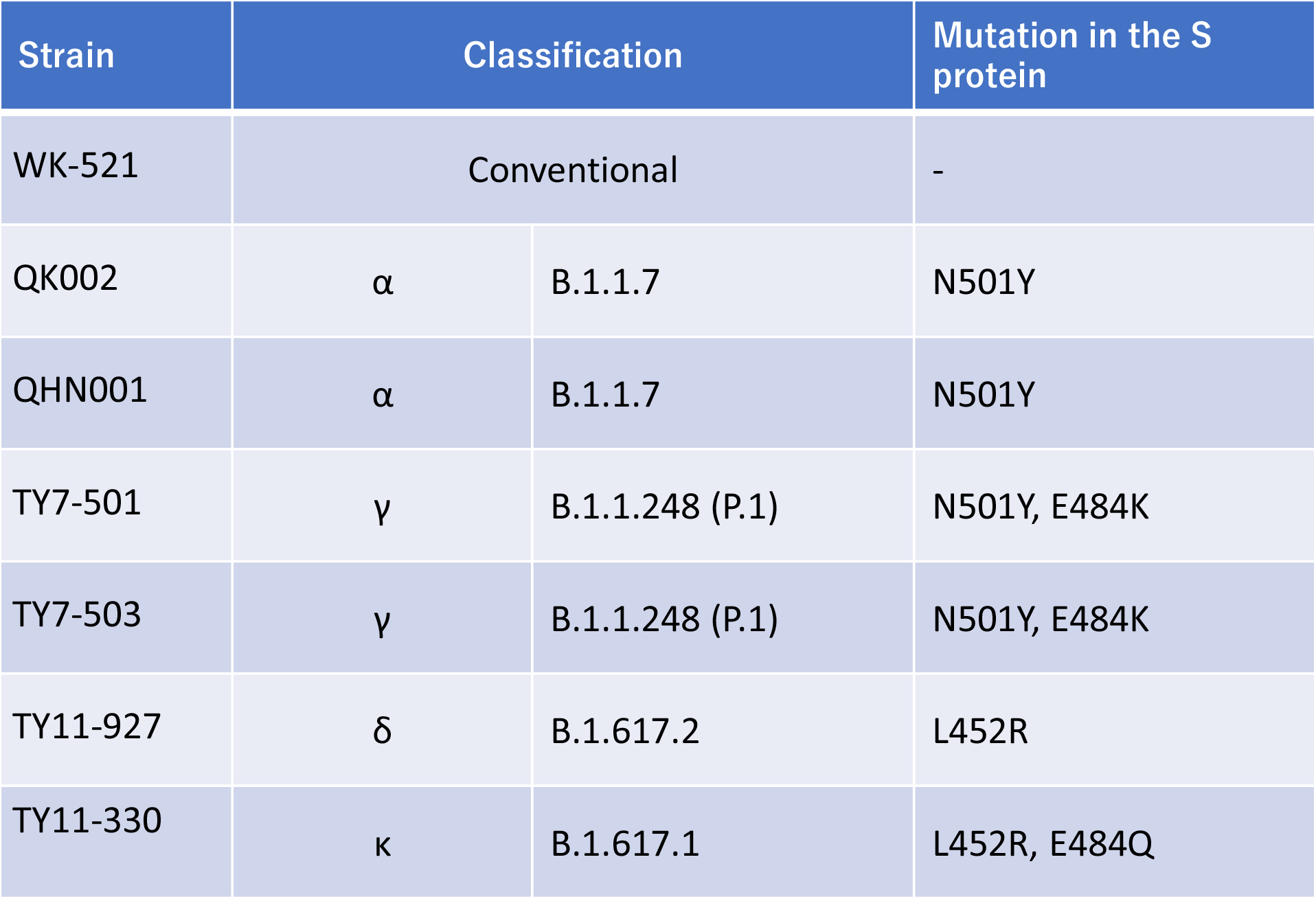
Viruses used in the study. All viruses were kindly provided by Japan National Institute of Infectious Diseases (Tokyo, Japan).

| Strain | Classification |  | Mutation in the S protein |
| --- | --- | --- | --- |
| WK-521 | Conventional |  | - |
| QK002 | $\alpha$ | B.1.1.7 | N501Y |
| QHN001 | $\alpha$ | B.1.1.7 | N501Y |
| TY7-501 | $\gamma$ | B.1.1.248 (P.1) | N501Y, E484K |
| TY7-503 | $\gamma$ | B.1.1.248 (P.1) | N501Y, E484K |
| TY11-927 | $\delta$ | B.1.617.2 | L452R |
| TY11-330 | $\kappa$ | B.1.617.1 | L452R, E484Q |

**Supplementary Table 2.** Donors of saliva used in the study. Saliva was purchased from Lee Biosolutions (Maryland Heights, MO, USA).

| Number | Birth year | Gender | Race |
| --- | --- | --- | --- |
| 1 | 1962 | Male | Caucasian |
| 2 | 1986 | Female | Caucasian |
| 3 | 1981 | Female | Caucasian |
| 4 | 1987 | Male | Caucasian |
| 5 | 1989 | Male | African American |
| 6 | 1977 | Female | Caucasian |
| 7 | 1967 | Female | Caucasian |
| 8 | 1991 | Male | Caucasian |

## References

1. Khan, S.; Liu, J.; Xue, M. Transmission of SARS-CoV-2, Required Developments in Research and Associated Public Health Concerns. Front Med (Lausanne) 2020, 7, 310, doi:10.3389/fmed.2020.00310.

2. Li, Y.; Ren, B.; Peng, X.; Hu, T.; Li, J.; Gong, T.; Tang, B.; Xu, X.; Zhou, X. Saliva is a non-negligible factor in the spread of COVID-19. Mol Oral Microbiol 2020, 35, 141–145, doi:10.1111/omi.12289.

3. Peng, X.; Xu, X.; Li, Y.; Cheng, L.; Zhou, X.; Ren, B. Transmission routes of 2019-nCoV and controls in dental practice. Int J Oral Sci 2020, 12, 9, doi:10.1038/s41368-020-0075-9.

4. Patel, K.P.; Vunnam, S.R.; Patel, P.A.; Krill, K.L.; Korbitz, P.M.; Gallagher, J.P.; Suh, J.E.; Vunnam, R.R. Transmission of SARS-CoV-2: an update of current literature. Eur J Clin Microbiol Infect Dis 2020, 39, 2005–2011, doi:10.1007/s10096-020-03961-1.

5. Wang, C.C.; Prather, K.A.; Sznitman, J.; Jimenez, J.L.; Lakdawala, S.S.; Tufekci, Z.; Marr, L.C. Airborne transmission of respiratory viruses. Science 2021, 373, doi:10.1126/science.abd9149.

6. Ohgitani, E.; Shin-Ya, M.; Ichitani, M.; Kobayashi, M.; Takihara, T.; Kawamoto, M.; Kinugasa, H.; Mazda, O. Rapid Inactivation In Vitro of SARS-CoV-2 in Saliva by Black Tea and Green Tea. Pathogens 2021, 10, doi:10.3390/pathogens10060721.

7. Ohgitani, E.; Shin-Ya, M.; Ichitani, M.; Kobayashi, M.; Takihara, T.; Kawamoto, M.; Kinugasa, H.; Mazda, O. Significant Inactivation of SARS-CoV-2 In Vitro by a Green Tea Catechin, a Catechin-Derivative, and Black Tea Galloylated Theaflavins. Molecules 2021, 26, doi:10.3390/molecules26123572.

8. Sanyaolu, A.; Okorie, C.; Marinkovic, A.; Haider, N.; Abbasi, A.F.; Jaferi, U.; Prakash, S.; Balendra, V. The emerging SARS-CoV-2 variants of concern. Ther Adv Infect Dis 2021, 8, 20499361211024372, doi:10.1177/20499361211024372.

9. Boehm, E.; Kronig, I.; Neher, R.A.; Eckerle, I.; Vetter, P.; Kaiser, L.; Geneva Centre for Emerging Viral, D. Novel SARS-CoV-2 variants: the pandemics within the pandemic. Clin Microbiol Infect 2021, 27, 1109–1117, doi:10.1016/j.cmi.2021.05.022.

10. Kannan, S.; Shaik Syed Ali, P.; Sheeza, A. Evolving biothreat of variant SARS-CoV-2 - molecular properties, virulence and epidemiology. Eur Rev Med Pharmacol Sci 2021, 25, 4405–4412, doi:10.26355/eurrev_202106_26151.

11. Aguie-Beghin, V.; Sausse, P.; Meudec, E.; Cheynier, V.; Douillard, R. Polyphenol-beta-casein complexes at the air/water interface and in solution: effects of polyphenol structure. J Agric Food Chem 2008, 56, 9600–9611, doi:10.1021/jf801672x.

12. Jobstl, E.; Howse, J.R.; Fairclough, J.P.; Williamson, M.P. Noncovalent cross-linking of casein by epigallocatechin gallate characterized by single molecule force microscopy. J Agric Food Chem 2006, 54, 4077–4081, doi:10.1021/jf053259f.

13. Bourassa, P.; Cote, R.; Hutchandani, S.; Samson, G.; Tajmir-Riahi, H.A. The effect of milk alpha-casein on the antioxidant activity of tea polyphenols. J Photochem Photobiol B 2013, 128, 43–49, doi:10.1016/j.jphotobiol.2013.07.021.

14. Haratifar, S.; Corredig, M. Interactions between tea catechins and casein micelles and their impact on renneting functionality. Food Chem 2014, 143, 27–32, doi:10.1016/j.foodchem.2013.07.092.

15. Holt, C.; Carver, J.A.; Ecroyd, H.; Thorn, D.C. Invited review: Caseins and the casein micelle: their biological functions, structures, and behavior in foods. J Dairy Sci 2013, 96, 6127–6146, doi:10.3168/jds.2013-6831.

16. Severin, S.; Wenshui, X. Milk biologically active components as nutraceuticals: review. Crit Rev Food Sci Nutr 2005, 45, 645–656, doi:10.1080/10408690490911756.

17. Pereira, P.C. Milk nutritional composition and its role in human health. Nutrition 2014, 30, 619–627, doi:10.1016/j.nut.2013.10.011.

18. Lorenz, M.; Jochmann, N.; von Krosigk, A.; Martus, P.; Baumann, G.; Stangl, K.; Stangl, V. Addition of milk prevents vascular protective effects of tea. Eur Heart J 2007, 28, 219–223, doi:10.1093/eurheartj/ehl442.

19. Rashidinejad, A.; Birch, E.J.; Sun-Waterhouse, D.; Everett, D.W. Addition of milk to tea infusions: Helpful or harmful? Evidence from in vitro and in vivo studies on antioxidant properties. Crit Rev Food Sci Nutr 2017, 57, 3188–3196, doi:10.1080/10408398.2015.1099515.

20. Tchesnokova, V.; Kulasekara, H.; Larson, L.; Bowers, V.; Rechkina, E.; Kisiela, D.; Sledneva, Y.; Choudhury, D.; Maslova, I.; Deng, K.; et al. Acquisition of the L452R mutation in the ACE2-binding interface of Spike protein triggers recent massive expansion of SARS-CoV-2 variants. J Clin Microbiol 2021, JCM0092121, doi:10.1128/JCM.00921-21.

21. Motozono, C.; Toyoda, M.; Zahradnik, J.; Saito, A.; Nasser, H.; Tan, T.S.; Ngare, I.; Kimura, I.; Uriu, K.; Kosugi, Y.; et al. SARS-CoV-2 spike L452R variant evades cellular immunity and increases infectivity. Cell Host Microbe 2021, 29, 1124–1136 e1111, doi:10.1016/j.chom.2021.06.006.

22. Huang, N.; Perez, P.; Kato, T.; Mikami, Y.; Okuda, K.; Gilmore, R.C.; Conde, C.D.; Gasmi, B.; Stein, S.; Beach, M.; et al. SARS-CoV-2 infection of the oral cavity and saliva. Nat Med 2021, doi:10.1038/s41591-021-01296-8.

23. Matsuyama, S.; Nao, N.; Shirato, K.; Kawase, M.; Saito, S.; Takayama, I.; Nagata, N.; Sekizuka, T.; Katoh, H.; Kato, F.; et al. Enhanced isolation of SARS-CoV-2 by TMPRSS2-expressing cells. Proc Natl Acad Sci U S A 2020, 117, 7001–7003, doi:10.1073/pnas.2002589117.

24. Pezzotti, G.; Ohgitani, E.; Shin-Ya, M.; Adachi, T.; Marin, E.; Boschetto, F.; Zhu, W.; Mazda, O. Instantaneous “catch-and-kill” inactivation of SARS-CoV-2 by nitride ceramics. Clin Transl Med 2020, 10, e212, doi:10.1002/ctm2.212.

